# The landscape of genomic structural variation in Indigenous Australians

**DOI:** 10.1101/2023.10.17.562810

**Authors:** Andre L.M. Reis, Melissa Rapadas, Jillian M. Hammond, Hasindu Gamaarachchi, Igor Stevanovski, Meutia Ayuputeri Kumaheri, Sanjog R. Chintalaphani, Duminda S.B. Dissanayake, Owen M. Siggs, Alex W. Hewitt, Bastien Llamas, Alex Brown, Gareth Baynam, Graham J. Mann, Azure Hermes, The National Centre for Indigenous Genomics, Hardip R. Patel, Ira W. Deveson

## Abstract

Indigenous Australians harbour rich and unique genomic diversity. However, Aboriginal and Torres Strait Islander ancestries are historically under-represented in genomics research and almost completely missing from reference databases. Addressing this representation gap is critical, both to advance our understanding of global human genomic diversity and as a prerequisite for ensuring equitable outcomes in genomic medicine. Here, we apply population-scale whole genome long-read sequencing to profile genomic structural variation across four remote Indigenous communities. We uncover an abundance of large indels (20-49bp; *n*=136,797) and structural variants (SVs; ≥50bp; *n*=159,912), the majority of which are composed of tandem repeat or interspersed mobile element sequences (90%) and have not been previously annotated (73%). A large fraction of SVs appear to be exclusive to Indigenous Australians (>30%) and the majority of these are found in only a single community, underscoring the need for broad and deep sampling to achieve a comprehensive catalogue of genomic structural variation across the Australian continent. Finally, we explore short-tandem repeats (STRs) throughout the genome to characterise allelic diversity at 50 known disease loci, uncover hundreds of novel repeat expansion sites within protein-coding genes, and identify unique patterns of diversity and constraint among STR sequences. Our study sheds new light on the dimensions, diversity and evolutionary trajectories of genomic structural variation within and beyond Australia.

## INTRODUCTION

The Australian continent has the longest history of continuous human occupation outside Africa. Australia is home to hundreds of Aboriginal nations or clans who inhabited all geographical regions throughout the continent, prospering in their diverse environments^1^. Over 250 distinct languages were spoken at the time of European invasion and ∼150 of these survive today^2^. Indigenous communities practise the world’s oldest surviving cultures. These are highly varied, but commonly emphasise the importance of kinship, ancestry, and relationships to the landscape and environment^1^.

While the remarkable cultural and linguistic diversity of Indigenous Australians is well documented, their rich and unique genomic diversity is relatively unexplored. Indigenous peoples have been historically under-represented in genomics research globally and Aboriginal ancestries are currently absent from leading international genomics resources, including the *1000 Genomes Project* and *gnomAD* reference databases^3,4^. Such resources are central to the interpretation, diagnosis and treatment of genetic disease but have reduced utility for communities without appropriate representation^5^. There is a pressing need to close this Indigenous representation gap to ensure equitable outcomes from genomic medicine in Australia^6,7^. Moreover, as one of the six inhabited continents on earth, the current lack of genomic data from Australia is a major gap in our understanding of global human genomic diversity.

The National Centre for Indigenous Genomics (NCIG)’s mission is to address this gap, by engaging Aboriginal and Torres Strait Islander communities in genomics research (https://ncig.anu.edu.au/). The NCIG has developed exemplary frameworks for Indigenous genomics that prioritise community leadership, participation and data sovereignty. The NCIG aims to develop relationships of deep trust with Indigenous communities, cataloguing their genomic diversity and conducting research in a manner that is sustainable, ethical, and not only beneficial to the partner communities but also aligned with their own ways of knowing.

In this study, we performed population-scale long-read sequencing using Oxford Nanopore Technologies (ONT) in four NCIG-partnered Aboriginal communities across northern and central Australia, as well as non-Indigenous Australians. The use of long-read sequencing technology, in combination with the recently completed telomere-to-telomere human reference genome (*T2T-chm13*)^8^, allows us to explore uncharted Aboriginal genomic diversity. Long-reads can resolve repetitive or non-unique genes/regions that are intractable with dominant short-read sequencing platforms^9^. Long-reads are also superior for the detection of structural variants (SVs), which account for the majority of the differences between the genomes of any two individuals and at least ∼25% of their deleterious alleles, yet are poorly understood due to technical/analytical limitations^9–11^.

Our study is among the first to deploy long-read sequencing at population-scale and the first to do so on a minority and/or Indigenous cohort^12^. We begin to describe the landscape of genomic structural variation in Indigenous Australians and establish frameworks for interpreting this variation in the context of genomic medicine.

## RESULTS

### Population-scale long-read sequencing in Aboriginal communities

To explore genomic structural variation in Indigenous Australians, we performed whole genome ONT sequencing on individuals from four remote Aboriginal communities with whom the NCIG has developed partnerships: Tiwi Islands (Wurrumiyanga, Pirlangimpi and Millikapiti communities; NCIG-P1), Galiwin’ku (NCIG-P2), Titjikala (NCIG-P3) and Yarrabah (NCIG-P4). These span a wide geographic, cultural and linguistic landscape (**Fig1a**). We sequenced 9-41 individuals from each community, 121 in total. We also sequenced 18 non-Indigenous Australian individuals of European ancestry for comparison, and two reference individuals of European ancestry from the Genome in a Bottle project for control purposes (HG001, HG002; **Supplementary Table 1**)^13^. High molecular weight DNA was extracted from saliva or blood and sequenced on an ONT PromethION device (see **Methods**). Across the complete cohort (*n* = 141), we obtained a median of ∼30-fold (14-47) genome coverage and ∼9.2 kb (2.7-16.8 kb) read-length N50 (**Extended Data Fig1a**,**1b**). Whilst DNA samples varied in quality, we obtained a minimum ∼10-fold coverage in reads of at least ∼5kb for every individual, providing a strong foundation for profiling genomic structural variation across the cohort (**Fig1b**; **Extended Data Fig1c**).

**Fig1.**
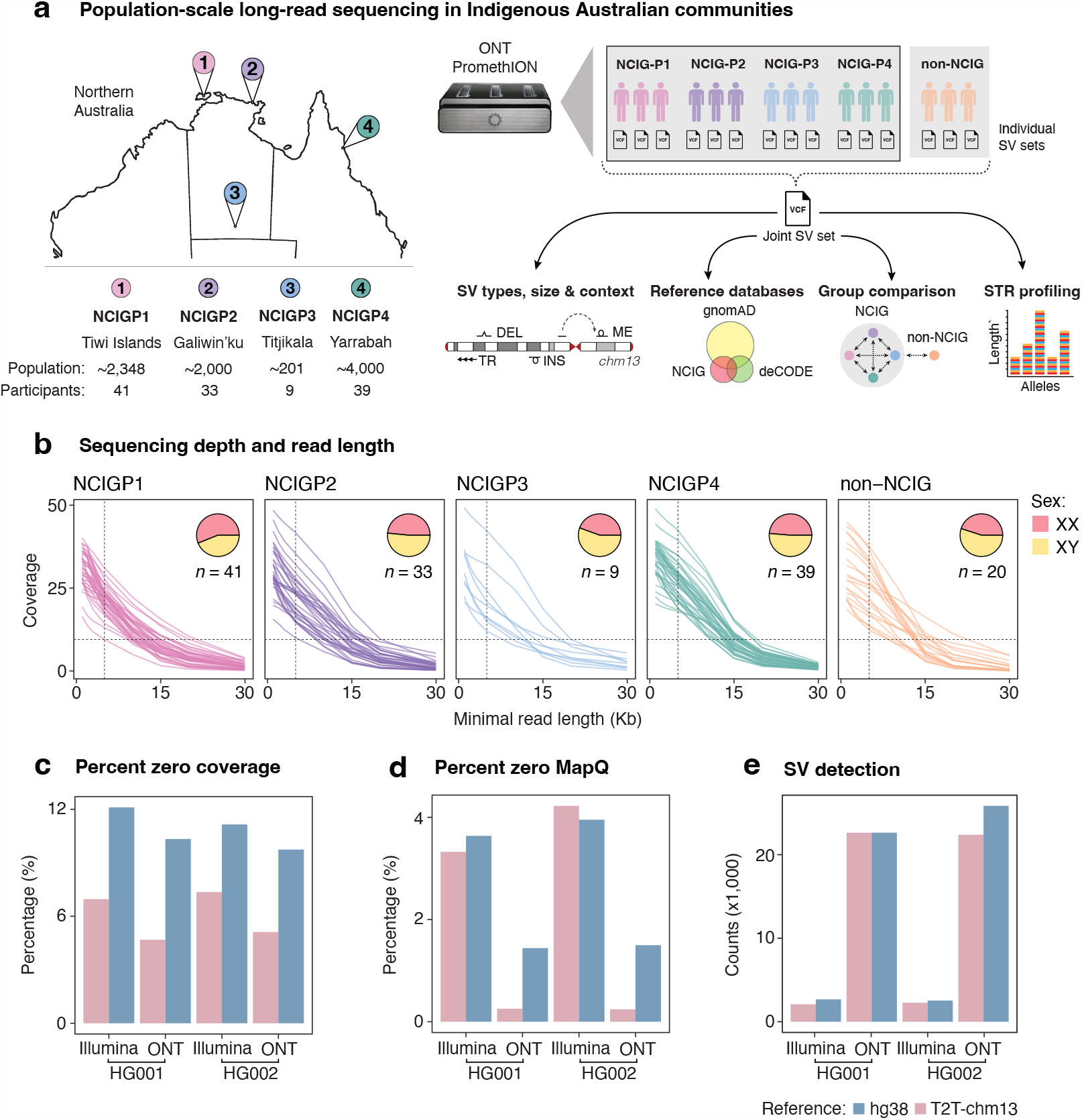
Long-read sequencing of Indigenous Australian communities. (**a**) Schematic illustrating the study design and analysis workflow. DNA samples were collected from four Indigenous communities in northern/central Australia: Tiwi Islands (NCIG-P1), Galiwin’ku (P2), Titjikala (P3), Yarrabah (P4), and from unrelated European individuals (non-NCIG). Map shows geographic locations, with population sizes and participant numbers underneath. Whole genome ONT sequencing was performed and reads were aligned to the T2T-chm13 genome. Structural variants (SVs) were called for each individual, then joint-calling performed to generate a non-redundant set of SVs, genotyped for each individual. SVs were characterised by type, size, context and compared to existing SV reference databases. SVs were compared between individuals and communities, with non-NCIG individuals as an outgroup. Short-tandem repeat (STR) alleles were genotyped to assess variation across communities. (**b**) Line plots show the average genomic coverage as sequencing reads were filtered by a minimum read-length cutoff. Each line represents one individual. Pie charts show the proportion of male (XY: yellow) and female (XX: red) participants from each community. (**c**) Percentage of genome with zero coverage for Illumina short-read and ONT long-read libraries from HG001 and HG002, when aligned to either hg38 (blue) or T2T-chm13 (pink). (**d**) Percentage of genome covered by alignments with low mapping quality (MAPQ<5) for the same comparisons. (**e**) Number of SVs detected with the same libraries and reference genomes as above.

Publication of the first complete human genome^8^ was a landmark for the field but, so far, there are few major studies outside the T2T Consortium that have used *T2T-chm13* as their chosen reference genome. We evaluated mappability and SV detection against the *T2T-chm13* reference, by comparison to *hg38*, using both ONT and short-read sequencing data from the HG001 and HG002 reference samples (see **Methods**). As expected, ONT data exhibited superior unique-alignment coverage and more comprehensive SV detection than short-reads (**Fig1c-e**). These advantages were further enhanced by use of the *T2T-chm13* reference, which had proportionally fewer regions of zero coverage (mean 4.9% vs 10.0%) or low mapability (MAPQ < 5; mean 0.2% vs 1.4%) and, as a result, afforded an additional ∼125 Mbases of total reference sequence that was accessible to analysis with ONT data (**Fig1c,d**). Manual inspection of medically relevant repetitive genes, such as *MUC1*^*14*^ (**Extended Data Fig2**), showed these were generally best resolved using the combination of ONT and *T2T-chm13*. Taken together, these results highlight the advantages of long-read sequencing and *T2T-chm13* for profiling genomic structural variation at high resolution.

Adopting *T2T-chm13* as our genomic reference, we called SVs (≥ 50bp) and large indels (20-49bp) in each individual (*CuteSV*^15^; **Fig1a**). Variants were filtered to exclude events with weak evidence (QUAL ≥ 5). We detected 21,723 SVs, on average, per individual, of which 19,089 were retained after filtering (10,126 deletions and 8,893 insertions; **Extended Data Fig1d**). The latter count is somewhat lower than reported in several recent long-read sequencing studies^16,17^, reflecting our preference for retaining only high-confidence SVs. Callsets were then merged (*Jasmine*^18^) into a unified joint-call catalogue comprising 159,912 thousand unique SVs and 136,797 thousand large indels, with which to explore the landscape of genomic structural variation in Indigenous Australians (**Fig1a**; see **Methods**). Notably, this surpassess the 134,886 SVs recently identified by ONT sequencing of 3,622 Icelanders^16^ (the largest long-read sequencing study to date), reflecting far greater genetic heterogeneity in our smaller cohort.

### Landscape of genomic structural variation

To better characterise the landscape, we next stratified variants (indels and SVs) by type, size and context (**Fig1a**; see **Methods**). A clear majority of all non-redundant variants (84.9%) were composed of repetitive sequences, including 103,425 short-tandem repeat (STR; 2-12bp) and 123,667 tandem-repeat (TR; >12bp) expansions/contractions, and 25,096 insertions/deletions of interspersed mobile element sequences (**Fig2a**). We detected 6,947 (2.3%) homopolymeric deletions/insertions, although we note these are likely to be enriched for technical errors, based on known ONT sequencing error profiles^19^. The remaining 37,574 variants (12.6%) were found to be non-repetitive (**Fig2a**).

**Fig2.**
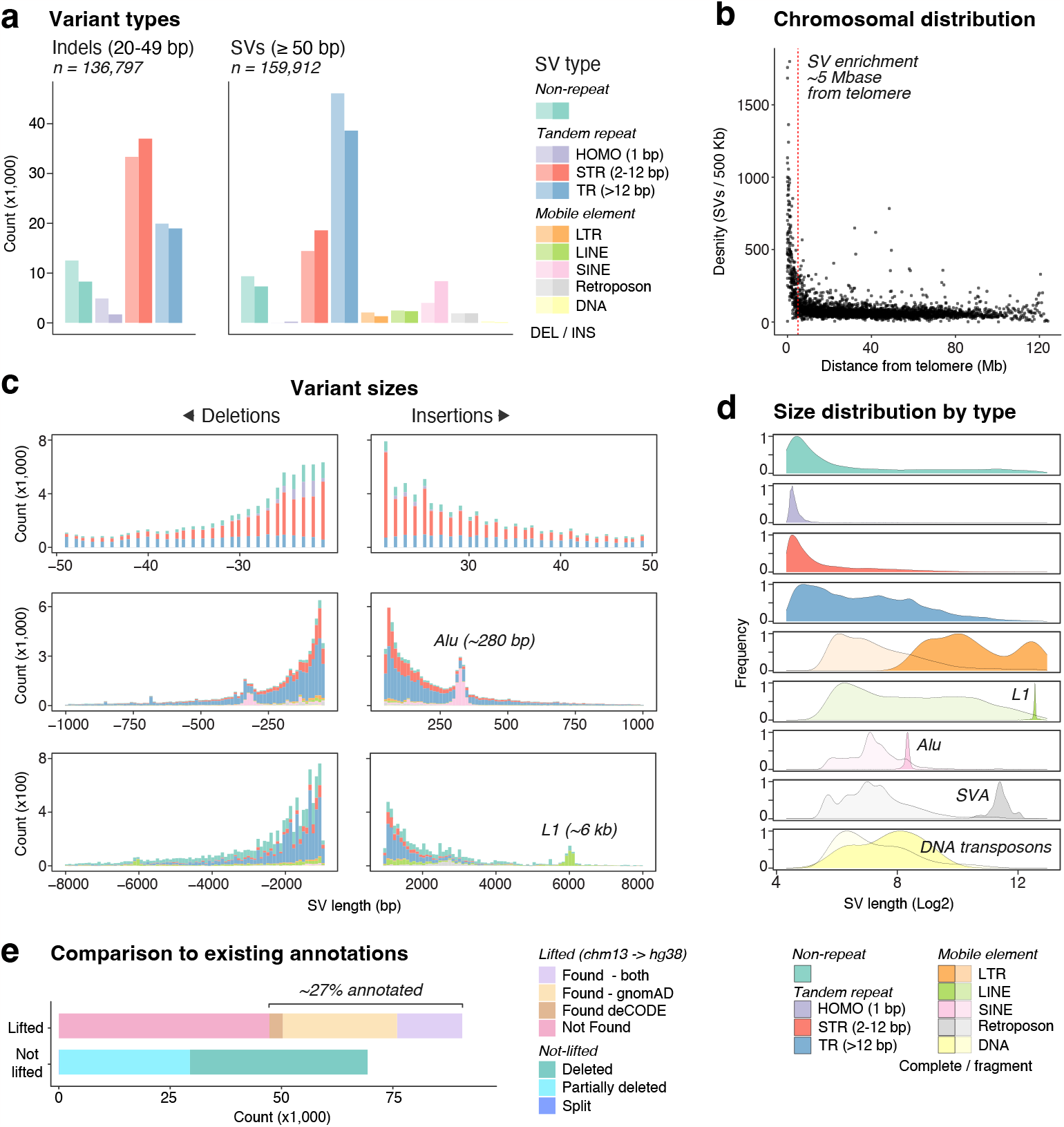
Landscape of genomic structural variation. (**a**) Number of non-redundant variants identified across the cohort (n = 141). Large indels (20-49bp; left) and structural variants (SVs; ≥50bp; right) are shown separately and parsed by type: non-repetitive (teal), tandem repeats (HOMO/homopolymers = purple, STR/short tandem repeats = red & TR/tandem repeats = blue) and mobile elements (LTR/long terminal repeat = orange, LINE/long interspersed nuclear elements = green, SINE/short interspersed nuclear element = pink, Retroposon = gray & DNA/DNA transposons = yellow). Light shades represent deletions and dark shades insertions. (**b**) Frequency of non-redundant SVs relative to distance from the nearest telomere, as calculated in 500 kb windows and averaged across all chromosomes. (**c**) Size distribution of insertions (positive values) and deletions (negative values), parsed by type (colour scheme as above). Three different resolution levels are displayed. Characteristic peaks for Alu elements (280 bp) and L1 elements (6 kb) are marked. (**d**) Size distributions for each variant type. Mobile element SVs are classified as ‘complete’ (dark shade) if they encompass one or more complete annotated element or ‘fragment’ (light shade), if only part. (**e**) Number of non-redundant SVs found in the gnomAD and/or deCODE reference databases. SVs were first lifted from T2T-chm13 to hg38. Some could not be lifted because they were deleted (green) or partially deleted (cyan) in hg38. SVs that could be lifted were classified as ‘found’ in gnomAD (beige), deCODE (brown), both (purple) or ‘not found’ in either (pink).

Structural variation was not distributed evenly across the genome, but showed higher density within ∼5Mbase of the telomere on each chromosome (**Fig2b**), as has been reported elsewhere^16,20^. This effect was almost entirely driven by TR-associated SVs, which were strongly enriched in sub-telomeric regions, with other classes being evenly distributed (**Extended Data Fig3a**). Both metacentric and acrocentric chromosomes were similarly affected (**Extended Data Fig3b**).

We observed characteristic differences in size between variants of different types (**Fig2c,d**). TR-associated SVs were generally larger than STR or non-repetitive SVs. Size distributions for mobile element SVs displayed clear peaks around expected sizes for prominent repeat families, including Alu (SINE; ∼280bp), L1 (LINE; ∼6kb) and SVA (Retroposon; ∼2kb; **Fig2c,d**)^21^. While most SVs associated with mobile elements encompassed only part of an annotated element, the aforementioned peaks are formed by SVs encompassing one or more complete element of a given type, which represent transposition events occurring since the common ancestor of individuals in our cohort and the *T2T-chm13* reference (European origin; **Fig2d**). Of these complete elements, SINEs were dominant (8,867), reflecting comparatively high Alu activity, and significant numbers of LINE (501) and Retroposon (327) transpositions were also detected (**Fig2a,d**). Although non-repetitive elements were predominantly short (**Fig2d**), the majority of very large SVs (>50kb, *n* = 117/121) were non-repetitive, including deletions detected up to ∼882 Kb in length.

Given the inclusion of unique, under-represented Australian communities, the use of population-scale long-read sequencing and the *T2T-chm13* reference, our catalogue contained a high proportion of SVs that have not been previously annotated (**Fig2e**). To assess novelty, we compared SVs to: (*i*) the *gnomAD* SV database^10^, which spans a diverse global cohort sequenced on short-read platforms; and (*ii*) an SV callset published recently by *deCODE* genetics^16^, based on population-scale ONT sequencing of Icelandic individuals. For this analysis, it was necessary to first convert SV coordinates to the *hg38* reference, on which these annotations are based (*LiftOver*). A significant number of SVs could not be lifted from *T2T-chm13* to *hg38* because their corresponding positions were fully (24.9%) or partially (18.3%) missing from the latter (**Fig2e**). Of the 90,578/159,912 SVs that were successfully lifted to *hg38*, we found a similar SV in either the *gnomAD* or *deCODE* catalogues for 43,289, or just ∼27% of all non-redundant SVs in the catalogue.

### Distribution and diversity

We next assessed the distribution of genomic structural variation among Indigenous and non-Indigenous individuals in the cohort. Overall, the majority of all non-redundant SVs were either private (i.e. found in a single individual; 26.3%) or polymorphic (<50% of individuals; 65.5%), with the remaining being classified as major (>50% of individuals; 7.8%) or shared alleles (all individuals; 0.2%; **Extended Data Fig4b**). Although different SV types were distributed uniformly among individuals and communities (**Fig3a**; **Extended Data Fig4a**), they varied in the degree to which they were shared between individuals (**Fig3b**; **Extended Data Fig4c**). For example, the proportional representation of STR- and TR-associated SVs was skewed towards polymorphic/private variation, whereas mobile elements and non-repetitive SVs were proportionally enriched among major/shared variation (**Fig3b**). These trends indicate the different rates at which different classes of SVs emerge and change over generations, presumably reflecting differences in their underlying mechanisms of formation.

**Fig3.**
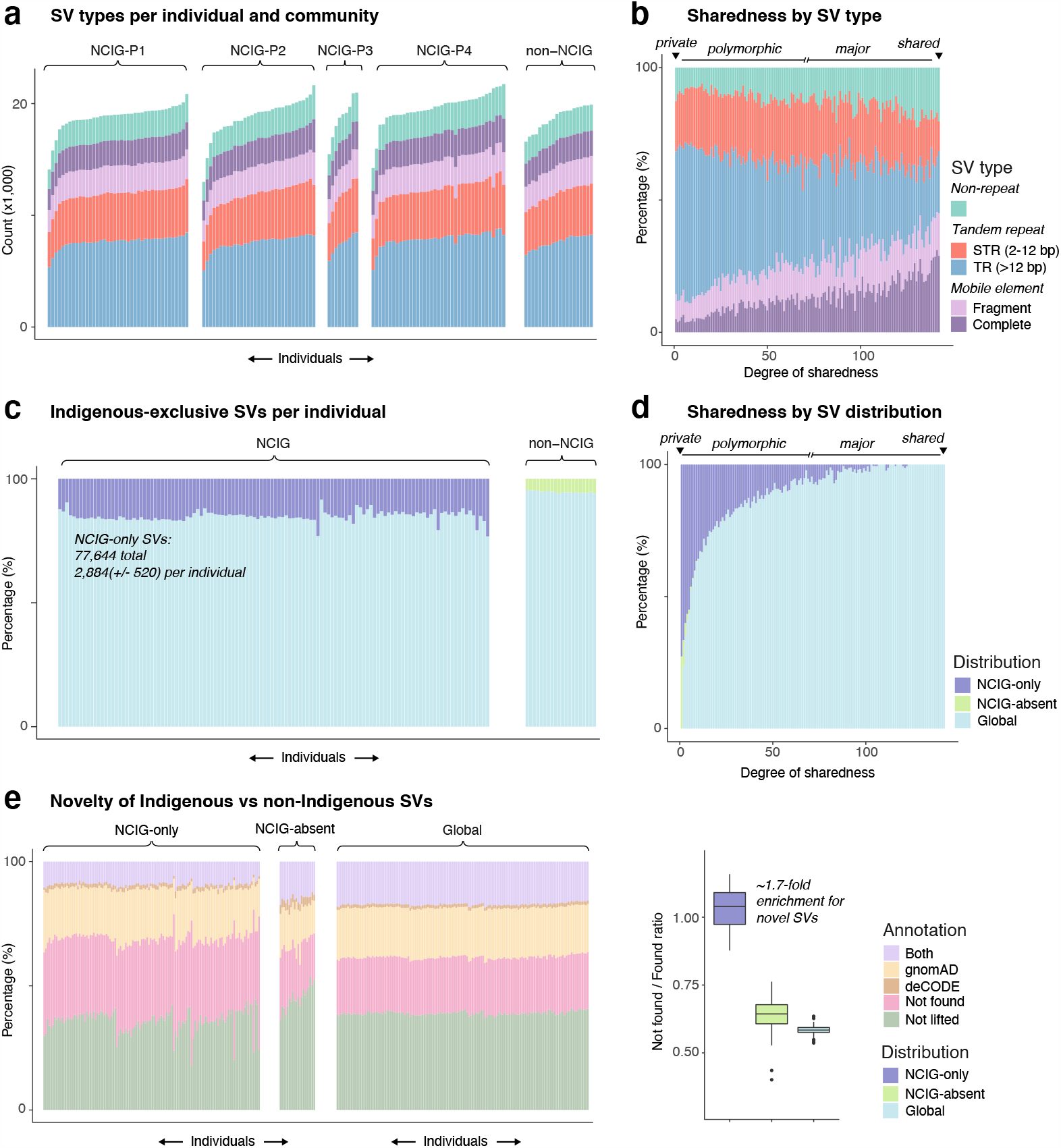
Distribution of SVs in Indigenous and non-Indigenous individuals. (**a**) Number of SVs identified in individuals from each group (NCIG-P1, P2, P3, P4 & non-NCIG), parsed by type: non-repetitive (teal), tandem repeat (STR = red & TR = blue) and mobile element (fragment = light purple & complete = dark purple). (**b**) Proportional representation of different types for SVs identified within a given number of individuals (‘degree of sharedness’). SVs were labelled as ‘private’ (1 individual), ‘polymorphic’ (≥ 2 & < 50% of individuals), ‘major’ (≥ 50% of individuals & < all individuals) and ‘shared’ (all individuals). (**c**) Proportion of SVs in each individual that were found exclusively in NCIG individuals (NCIG-only = purple), or exclusively in non-NCIG individuals (NCIG-absent = green), or across both (Global = light blue). (**d**) Proportional representation of NCIG-only, NCIG-absent and Global SVs according to degree of sharedness (as above). (**e**) Proportion of NCIG-only, NCIG-absent or Global SVs in each individual that were previously annotated in the gnomAD or deCODE databases (see **Fig2e**). Accompanying boxplot shows the distribution of ratios between the proportion of SVs ‘not found’ in either gnomAD or deCODE, to the proportion of SVs ‘found’ in one or both annotations. All panels in Fig3 display results for SVs (≥ 50bp) only; equivalent plots for large indels (20-49bp) are shown in **Extended Data Fig4**.

A large proportion of SVs across the complete non-redundant catalogue was seen only among Indigenous individuals (‘NCIG-only’; 48.5%) or only among non-Indigenous participants (‘NCIG-absent’; 9.2%; **Fig3c**). NCIG-only SVs made up a significantly higher proportion of total SVs in a given Indigenous individual (15.0% ± 2.0) than NCIG-absent SVs in non-Indigenous individuals (5.2% ± 0.4; **Fig3c**; **Extended Data Fig4d**). The majority of NCIG-only variants were polymorphic (**Fig3d**; **Extended Data Fig4e**) and were previously unannotated – more so than for NCIG-absent variants (**Fig3e**). On average, each Indigenous individual harboured ∼2,884 ± 520 NCIG-only SVs, of which ∼965 ± 328 were unannotated and may therefore represent exclusively Australian Indigenous variation.

Principle coordinate analysis (PCoA) of structural variation further reiterated the clear genetic distinctions between Indigenous and non-Indigenous individuals (**Fig4a**). This also highlighted the distinct genetic architecture of different communities, which formed largely separate PCoA clusters (**Fig4a**). Indeed, among Indigenous individuals, we found that 56.4% of NCIG-only SVs were found in just a single individual or community, whereas NCIG-only SVs shared between more than one individual across all communities were relatively rare (2.8%; **Fig4b**; **Extended Data Fig5a**). Shared SVs showed a proportional enrichment of mobile elements and a depletion of TR SVs (**Extended Data Fig5b**), consistent with the contrasting polymorphism for these SV types (**Fig3b**). Of the ∼965 exclusively Indigenous SVs in a given individual identified above, an average of ∼593 ± 132 were not found outside their community.

**Fig4.**
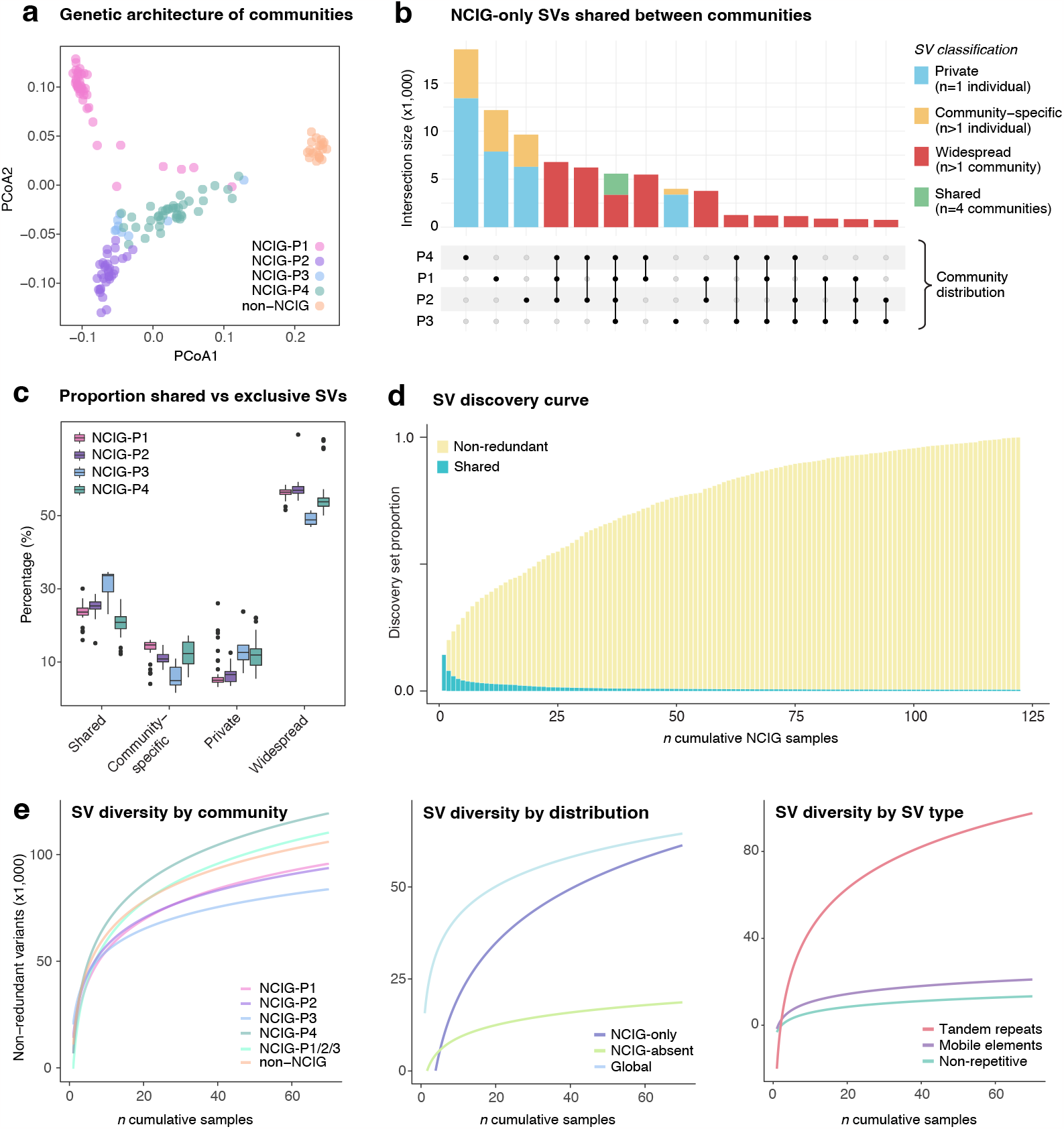
Distribution of SVs between Indigenous communities. (**a**) Principal coordinate analysis (PCoA) representing the distance between individuals in the cohort based on their SV compositions. Each dot represents an individual, coloured according to their group (NCIG-P1 = pink, P2 = purple, P3 = blue, P4 = green & non-NCIG = orange). (**b**) Distribution of ‘NCIG-only’ variants (see **Fig3**) shared among the four NCIG communities. SVs were classified as private (n = 1 individual; blue), community-specific (n > 1 individual in 1 community; yellow), widespread (n > 1 individual in more than 1 community; red) or shared (n > 1 individual in all 4 communities; green). (**c**) Proportion of private, community-specific, widespread & shared NCIG-only variants among individuals, grouped by community. (**d**) SV discovery curve in which, starting with a single NCIG individual, the number of new non-redundant SVs is counted as new individuals are iteratively added. SVs shared among all previously added samples are shown as green portions of each bar. (**e**) Log regression models predicting the number of non-redundant SVs identified, given the number of individuals sampled. The models are broken down by community (left panel), by geographical distribution (centre panel) and SV type (NCIG individuals; right panel). All panels in **Fig4** display results for SVs (≥ 50bp) only; equivalent plots for large indels (20-49bp) are shown in Extended Data Fig5.

Next, we generated discovery curves that model the diversity of structural variation within a set of individuals (see **Methods**). Across the 121 Indigenous individuals in the cohort, cumulative SV discovery did not approach saturation, indicating many further SV alleles remain to be sampled (**Fig4d**; **Extended Data Fig5d**). This diversity was not shared equally among different communities, nor variant types (**Fig4c-e**; **Extended Data Fig5c**). Individually, the Tiwi Islands (NCIG-P1), Galiwin’ku (P2) and Titjikala (P3) communities each showed lower within-community SV diversity than seen among the non-NCIG comparison group, reflecting their small population sizes and relative isolation – see **Fig1a**. In contrast, Yarrabah (P4) harboured substantially higher genomic diversity than the other communities, a higher proportion of private variation, and alone showed greater diversity than the non-Indigenous group (**Fig4c,e**). NCIG-only SVs showed greater heterogeneity among NCIG-individuals than for NCIG-absent SVs among non-NCIG individuals (**Fig4e**). Finally, we found that SV diversity was driven most strongly by TR-associated SVs, whereas new discovery of mobile elements and non-repetitive SVs was largely saturated (**Fig4e**; **Extended Data Fig5e**).

### Functional context

Given the vast diversity of genomic structural variation described above and the predominance of SV classes that are poorly studied, we next used measures of purifying selection to investigate their functional relevance. A large indel or SV intersecting with one or more CDS exon/s in a protein-coding gene is likely to truncate or alter its ORF, while a variant within an intron, UTR or proximal gene-regulatory region may impact transcription, translation or splicing. The extent to which these events disrupt gene function should be modelled by depletion of structural variation within essential genes among otherwise healthy populations^10^.

Across our complete cohort (*n* = 141), we detected 126,473 non-redundant variants (58,079 indels and 68,394 SVs) intersecting protein-coding loci, including 1,462 impacting CDS exons. An average individual possessed ∼156 ± 15 CDS variants and ∼20,124 ± 1,308 within non-CDS regions of protein-coding genes (introns, UTRs and proximal regulatory regions; **Extended Data Fig6a**). There was an enrichment of private variants intersecting CDS regions (33.7%) compared to the proportion of private variants (24.1%) in intergenic regions, consistent with purifying selection. Interestingly, variants intersecting CDS exons were almost all either non-repetitive (33.7%) or TR-associated (59.5%), with a strong depletion of STR and mobile element SVs in CDS, relative to intronic and intergenic regions (**Fig5a**).

**Fig5.**
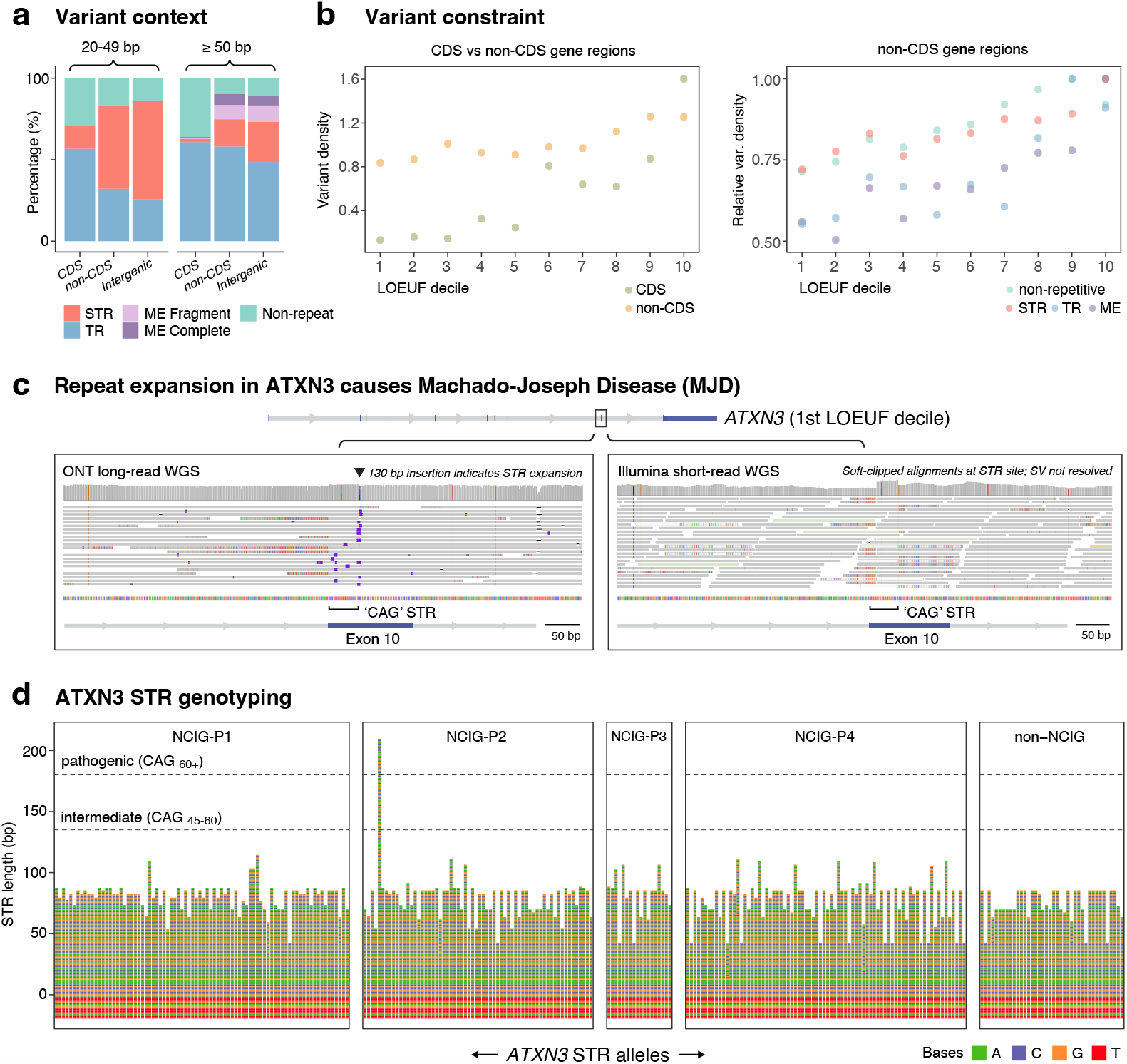
Functional relevance of genomic structural variation. (**a**) Proportion of different variant types identified within CDS exon/s, non-CDS regions of protein-coding genes (introns, UTRs and ± 2kb proximal regulatory regions) and intergenic regions, for large indels (20-49 bp; left) and SVs (≥ 50bp; right). Variants are classified as: non-repetitive (teal), tandem repeat (STR = red & TR = blue) and mobile element (fragment = light purple & complete = dark purple). (**b**; left) Variant density in CDS (green) and non-CDS (yellow) regions of protein-coding genes in different LOEUF deciles, which quantify intolerance to loss-of-function variation (1st decile = highest constraint). (**b**; right) As above, but non-CDS regions are parsed by variant type and normalised to their 10th decile. (**c**) Genome browser view shows sequencing alignments to *ATXN3*. A ‘CAG’ STR expansion, known to cause Machado-Joseph Disease (MJD), was identified in one NCIG-P2 individual. ONT reads span the expansion (left panel; purple markers indicate insertions). Illumina short-reads do not span the expansion, and are soft-clipped (right panel). (**d**) Sequence bar chart shows *ATXN3* STR alleles, including 20 bp of upstream flanking sequence, genotyped for every individual (two alleles each). Dashed lines show STR size cut-offs for intermediate and fully pathogenic alleles. A single pathogenic allele is identified in an NCIG-P2 individual. All others are in the ‘normal’ range, despite their diverse sizes. Nucleotide sequences are coloured, as follows: A (green), C (blue), G (orange) & T (red).

We next parsed protein-coding genes according to LOEUF, a metric that quantifies their intolerance to loss-of-function variation, developed previously by *gnomAD*^4^. This approach revealed clear constraint on structural variation in CDS regions (**Fig5b**). For example, we saw a ∼12-fold reduction in the size-normalised density of structural variation among the most essential genes (LOEUF decile 1; 1.29 × 10^−5^) compared to the least essential genes (LOEUF decile 10; 1.60 × 10^−4^; **Fig5b**; see **Methods**). We also observed significant, albeit weaker, constraint on non-CDS variants, with a ∼1.5-fold difference in variant frequency between the highest and lowest deciles (**Fig5b**). This was not shared equally by SVs of different types; mobile element-associated SVs showed the strongest purifying selection (1.8-fold difference between highest/lowest deciles) and STR-associated SVs showed the weakest (1.3-fold; **Fig5b**). Assessing LOEUF among Indigenous-specific variation, we found that most SVs (81.1%) in CDS regions of essential genes were private or community-specific (**Extended Data Fig6b**).

Since evidence of selection is critical for interpreting the functional relevance of genetic variation, the findings above help to establish the suitability of our analysis framework and SV catalogue to inform genomic medicine applications in Indigenous Australians. Even among just 141 individuals sequenced here, we identified 69 deletions/insertions impacting CDS regions of genes in LOEUF decile 1-2, 44 of which were not previously annotated. This includes complete or near-complete deletions of essential genes including *PRKRA, BRD9, SHOX, NID2, PABPC5, JAZF1, NIPA2* and *ANOS1*.

One notable SV in an essential gene (LOEUF decile 1) was a 130 bp ‘CAG’ STR expansion in *ATXN3* that is known to cause Machado-Joseph Disease (MJD; a.k.a. Spinocerebellar ataxia type 3; SCA3)^22^. This pathogenic allele was detected in a single individual from Galiwin’ku community (NCIG-P2), with long-read sequencing clearly defining the size, position and sequence of the STR expansion, unlike short-read whole genome sequencing on the same individual (**Fig5c,d**). MJD is a late onset, progressive movement disorder with autosomal dominant inheritance and complete penetrance for expansions of this size^22^. MJD affects up to ∼5/100,000 people worldwide, but is estimated to be >100 times more prevalent among Indigenous populations in some areas of Australia’s Northern Territory (NT)^23^.

The individual in question had consented to receive reportable incidental genetic findings arising from NCIG research. This prompted an ongoing dialogue between NCIG, Galiwin’ku representatives, a local genetic counsellor and the MJD foundation (https://mjd.org.au/), who work with remote NT Aboriginal communities to develop unique clinical genetics service models tailored for their needs^24^. Under recommendation of the MJD Foundation, the genetic counsellor was able to contact the individual and their family, arranging for clinical testing and appropriate follow-up.

### Characterising short-tandem repeat expansions

The preceding example highlights the utility of long-read sequencing for profiling variation in STR sequences – both normal and pathogenic – as well as the importance of ancestry in interpreting this variation. STRs are highly polymorphic, and STR expansions are causative pathogenic variants in at least 37 neurogenetic and 10 congenital disorders^25^. However, STRs are refractory to analysis with short-read sequencing and, as a result, have been relatively poorly characterised to date, particularly among minority communities, such as Indigenous Australians. Our dataset provides a unique opportunity to explore allelic diversity in STR sequences at high-resolution and population-scale.

Across the cohort, we detected 55,595 non-redundant variants that constitute STR expansions (i.e., insertions) and 47,830 STR contractions (i.e., deletions; see **Methods**) relative to the *T2T-chm13* reference. These ranged in size from ∼20bp (our lower cutoff) to 99,204 bp and occurred predominantly in intergenic (63.9%) or non-CDS gene regions (35.9%; **Fig6a**). STR period size was negatively correlated with the global frequency of STR expansions/contractions. However, trinucleotide and hexanucleotide repeats were outliers from this trend, showing markedly lower frequencies than other periods (**Fig6a**). The opposite was true within CDS regions, where in-frame expansions (i.e. 3bp, 6bp, etc) occurred at higher frequencies than other periods (**Fig6a**). Therefore, although in-frame expansions/contractions are more tolerated within coding sequences (because they do not cause frameshifts), there appears to be higher constraint on in-frame STRs across the remainder of the genome. We hypothesise that this acts to limit the potential for spurious expression of toxic homomeric polypeptides that contribute to pathogenicity in many STR disorders^25^.

**Fig6.**
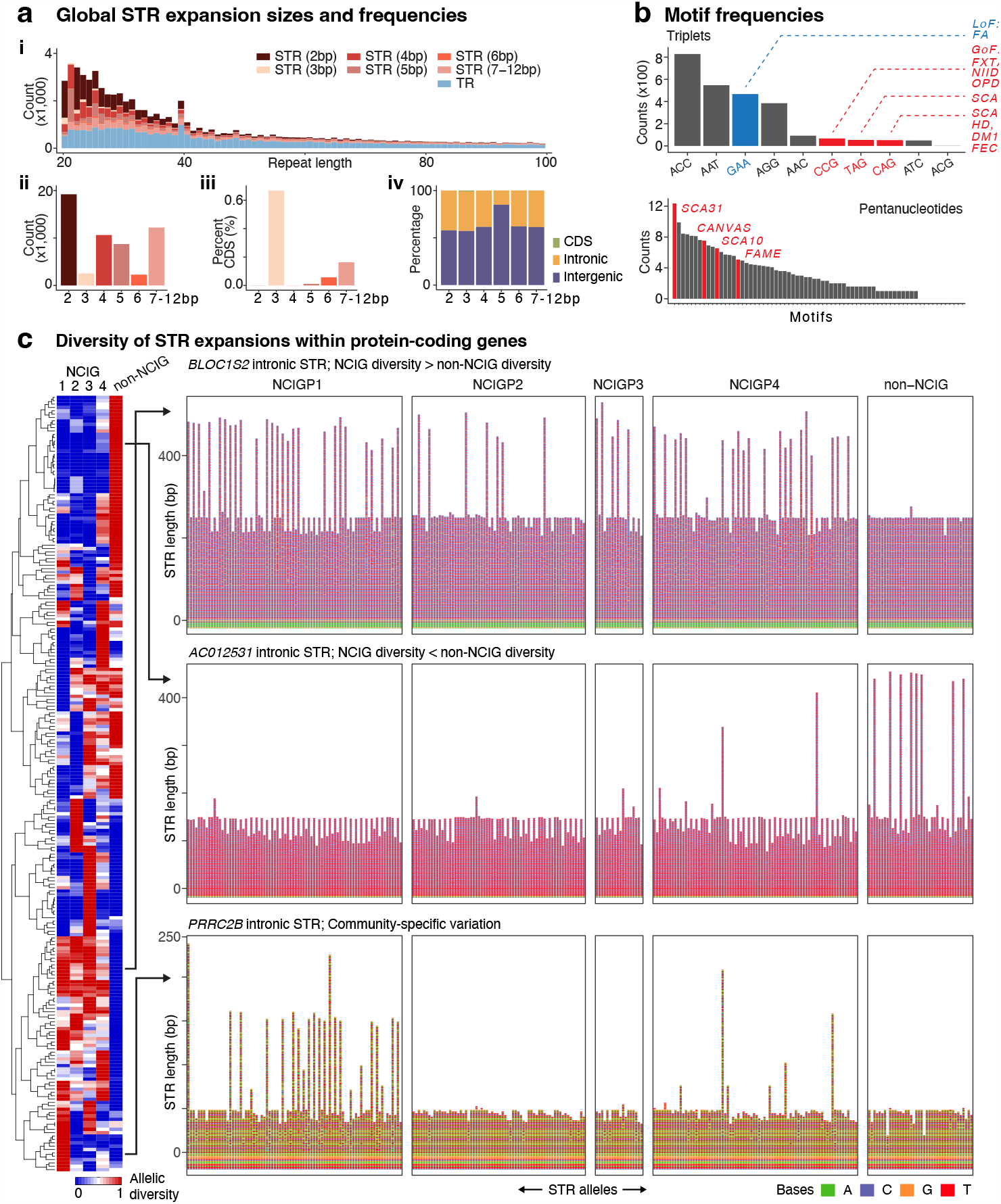
Landscape of STR expansions. (**ai**) Length distribution of all non-redundant tandem repeat (TR) and short tandem repeat (STR) insertions (i.e. expansions) detected across the cohort (n=141), broken down by period size. (**aii**) Number non-redundant STR expansions in each period. (**aiii**) Relative frequency of each period within CDS exons. (**aiv**) Proportional composition of each period based on genomic context (CDS = green, non-CDS = yellow & intergenic = dark blue). (**b**) Frequency of every possible triplet (upper) and pentanucleotide (lower) motif among non-redundant STR expansions. Motifs known to cause different repeat disorders are highlighted. For triplet disorders, pathogenic motifs with gain-of-function (GoF) mechanisms are in red, loss-of-function (LoF) in blue. (**c**; left) Normalised standard deviation (range 0-1) of allele sizes observed within each community group (matrix columns) for different STR sites (matrix rows). All expanded sites of period ≥3bp within protein-coding genes, in which allelic composition was significantly different between groups are shown. Hierarchical clustering groups STR sites based on patterns of variability between groups. (**c**; right) All STR alleles (two per individual) for three example sites showing distinct patterns of variation. *BLOC1S2* (top) has an intronic STR with higher allelic diversity in NCIG vs non-NCIG individuals. *AC012531* (middle) has lower allelic diversity in NCIG vs non-NCIG individuals. *PRRC2B* (bottom) shows community specific patterns, with NCIG-P1 and P4 exhibiting heterogeneity. Nucleotide sequences are coloured, as follows: A (green), C (blue), G (orange) & T (red).

Consistent with this idea, expansions of the STR motifs associated with poly-glutamine (‘CAG’) and poly-glicine (‘CGG’) disorders, which include MJD and a range of other disorders, were globally rare by comparison to other motifs (**Fig6b**)^25^. In contrast to these dominant, gain-of-function disorders, the ‘GAA’ expansion motif that triggers epigenetic silencing of *FXN* (i.e. loss-of-function) in Fredrich’s ataxia (FA), was relatively common, suggesting expansions of this motif are not typically deleterious in other contexts (**Fig6b**)^25^.

Besides triplets, intronic pentanucleotide STRs are most widely associated with known disorders — most of them described only recently^25^. In this context, it was notable that pentanucleotide STR expansions occurred at lower frequency (∼17%) within intronic regions than for other period sizes (∼40%), indicative of context-specific constraint (**Fig6a**). However, in contrast to triplet disorders, the motifs known to be associated with disease, such as ‘TGGAA’ (SCA31), ‘AAGGG’ (CANVAS) and ‘TTTCA’ (FAME) were among the most frequent pentanucleotide expansions (**Fig6b**). This may have implications for the mechanisms of pathogenicity in these disorders, which are currently not well understood.

While the SV joint-calling approach used throughout the preceding sections can identify expansion/contraction events, the process tends to collapse STR alleles of various sizes into a single consensus variant, rather than representing the full spectrum of alleles at a given site. To better resolve the STR landscape, we developed a method for individual-level, diploid STR genotyping, expansion discovery and visualisation (see **Methods**). We focused our analysis on 685 STR sites of period ≥3bp within protein-coding loci (including 7 within CDS exons) that were significantly expanded in at least one individual (**Extended Data Fig7a**), as well as 50 known disease-associated STR loci^25^. Using this approach, we stratified STR sites according to allelic diversity across communities, identifying 231 sites (18.9%) that showed inter-community differences in allelic composition (**Fig6c**; **Extended Data Fig7b**). We found 155 sites that were more diverse in Indigenous than non-Indigenous individuals and 76 sites where the opposite was true (**Fig6c**). Many STRs showed more local effects, such as elevated allelic diversity in just a single community, or expansions that were limited to a single, or a small handful, of individuals (**Fig6c**).

## DISCUSSION

Our study is among the first major genomic surveys of Indigenous Australians. Previous publications that include genomic data from Aboriginal and/or Torres Strait Islander peoples have largely focused on historical demographic processes^26–30^. These relied on mitochondrial^29,31^ or short-read whole genome sequencing^26–28,30^. Hence, we have generated the first suitable dataset to begin exploring the landscape of Indigenous genomic structural variation.

We found a diversity of structural variation across four remote Aboriginal communities in northern and central Australia. This was predominantly repetitive, encompassing thousands of tandem repeats and mobile elements. A significant proportion was only found in Indigenous individuals in our study, and have not been previously annotated in diverse global reference data^10,16^. However, exclusively Indigenous variants were generally not shared throughout the continent. The following picture emerges of structural variation in a given individual in the Indigenous communities studied here: >10 Mbases of the genome is affected by ∼28,000 large indels and ∼19,000 SVs; ∼950 SVs are not found outside Indigenous Australians; >60% of those Indigenous variants are not found in other (geographically separated) communities.

Our study sheds new light on the rich and unique genetic diversity of Indigenous Australians. Owing to the long history of continuous occupation, Australia’s Indigenous peoples are highly genetically distinct from non-Australians. This underscores the need for ancestry-appropriate reference data for genomic medicine, which is sorely lacking^5,6,7^. Moreover, Indigenous Australians should not be viewed as genetically homogenous. We show that different communities, clans and/or nations have highly distinct genomic architectures, mirroring their cultural and linguistic diversity^1^. Therefore, broad engagement — far beyond the four communities we have profiled here — will be required to adequately survey Indigenous genomic diversity and, ultimately, to achieve equitable outcomes in genomic medicine.

Our study is among a handful of efforts, globally, to implement long-read sequencing at population scale, and others have so far focused on comparatively homogeneous, well-studied populations (specifically Icelandic^16^, Chinese^32^ and Japanese^33^). Although our cohort was smaller than each of these studies, we identified a greater number of total non-redundant SVs, reflecting higher genetic heterogeneity among our participants. We use this rich catalogue to articulate a number of fundamental insights into the landscape of genomic structural variation in human populations that reach beyond Australia. For example, we show that: (*i*) SVs are predominantly repetitive, with TRs, STRs and mobile elements underpinning ∼87% of SVs per individual; (*ii*) SVs of different types and sizes show clear differences in their dynamics of inheritance, with TR and STR SVs being more polymorphic than mobile elements and non-repetitive SVs; (*iii*) SVs in both CDS and non-CDS regions of protein-coding genes are under purifying selection and different SV types show different signatures of constraint.

Our high-resolution survey of allelic diversity among STRs is similarly informative. We uncover an abundance of STR variation across the genome, including hundreds of previously undescribed STR expansions within protein-coding genes. STRs show distinct, context-specific signatures of selection; specific periods and motifs showing elevated constraint, globally. For both novel expansion sites and known disease-associated loci, we show pervasive Indigenous vs non-Indigenous and inter-community differences in STR allele composition. Constructing a clear picture of this complex background of normal STR variation is critical for the discovery and diagnosis of STR expansion disorders^34^. However, most existing reference data is based on European and East-Asian cohorts and, therefore, of reduced suitability for Indigenous Australian communities. Tangible examples of local effects are provided by a unique STR motif that causes CANVAS in individuals of Māori descent^35^ and MJD, an expansion disorder with dramatically elevated frequency in NT Aboriginal communities^23^. Our study begins to establish the appropriate context for interpreting Indigenous STR variation in future genomic medicine initiatives. Indeed, the incidental identification of a pathogenic MJD expansion in one individual highlights the strength of our approach in this domain.

## METHODS

### Cohorts

Saliva and/or blood samples were collected from consenting individuals among four NCIG-partnered communities: Tiwi Islands (comprising the Wurrumiyanga, Pirlangimpi and Millikapiti communities), Galiwin’ku, Titjikala and Yarrabah, between 2015 and 2019 (Ethics protocol number 2015/065). Non-Indigenous comparison data, generated from unrelated Australian individuals of European ancestry, was drawn from two existing biomedical research cohorts: (*i*) the Tasmanian Ophthalmic BioBank (Ethics protocol number 2020/ETH02479); and (*ii*) the Australian and New Zealand Registry of Advanced Glaucoma (Southern Adelaide Clinical Human Research Ethics Committee approval 305-08).

### Saliva sample collection and DNA extraction

Saliva samples were collected in Oragene-DNA collection tubes (OG-500, DNA Genotek Inc., Canada). Individuals were requested to avoid food intake 30 minutes prior to the collection and were asked to fill the collection tube to the best of their capacity. Approximately 3mL of total material (including stabilising liquid) was collected from individuals. Samples were transported to the NCIG lab in checked-in baggage in flight at the end of community visits. Saliva tubes were subject to large changes in temperature during collection and transport, and were kept at room temperature or at 4°C until further processing. Samples were split into 2 or more aliquots of 1mL each depending on the quantity of material available after heating samples at 50°C for 2 hours, as recommended by the manufacturer. Each sample tube was separately processed in a hood to reduce handling errors, cross contamination and external contamination. One of the aliquots with 1mL sample was used for the DNA extraction and remaining aliquots were stored at -20°C or -80°C for long term storage.

DNA extractions from saliva were performed by Australian Phenomics Facility (APF) on QIAsymphony SP using QIAsymphony DSP DNA Midi Kit (QIAGEN). Briefly, tubes were incubated at 56°C for 1 hour followed by addition of 2ul of RNAse A (100mg/ml, QIAGEN) to 1ml of saliva sample and incubated at room temperature for 5 minutes. RNAse activity was suppressed by incubation at 50°C for 40 minutes. A custom protocol on QIAsymphony SP, specifically developed for 1ml of saliva sample was then used to run the DNA extraction process on the instrument. DNA was eluted in 100uL of TE buffer.

### Fresh blood collection and DNA extraction

We collected fresh blood where possible from consenting individuals in BD Vacutainer® EDTA tubes (lavender caps, BD). Blood tubes were immediately placed on ice after the collection and shipped to the NCIG laboratory on dry ice. Blood samples were stored at -80°C until required. DNA extraction was performed using FlexiGene DNA Kit (QIAGEN) according to the manufacturer’s protocol. Briefly, frozen blood samples were thawed in 37°C water bath and mixed with the lysis buffer. A cell pellet was then collected by centrifugation at 2000g for 5 minutes and supernatant was discarded. Cell pellet was mixed with denaturation buffer and protease enzyme followed by incubation at 65°C in water bath for 10 minutes. DNA was precipitated using isopropanol and centrifuged at 2000g for 3 minutes. DNA pellet was then washed with 70% ethanol and pelleted at 2000g for 3 minutes. Finally, supernatant was discarded and the DNA pellet air dried. Dry DNA pellet was resuspended in 1mL of hydration buffer (10mM Tris-Cl) and incubated at 65°C for 1 hour for dissolution.

For non-Indigenous samples, HMW DNA was previously extracted from blood, using Qiagen DNeasy Blood and Tissue Kit, as per manufacturer’s instructions, and stored at -80°C.

### Whole genome ONT sequencing

HMW DNA samples were transferred to the Garvan Institute Sequencing Platform for long-read sequencing analysis on Oxford Nanopore Technologies (ONT) instruments. DNA quantity was measured using a Qubit (Thermo Fisher Scientific), purity on a NanoDrop (Thermo Fisher Scientific) and fragment-size distribution on a TapeStation (Agilent). Prior to ONT library preparations, DNA was sheared to ∼15-20kb fragment size using Covaris G-tubes. No shearing was performed on samples where the starting fragment distribution peaked at or below ∼25kb. Sequencing libraries were prepared from ∼1-2ug of DNA, using native library preparation kits (either SQK-LSK110 or SQK-LSK114), according to the manufacturer’s instructions. Each library was loaded onto a PromethION flow cell (R9.4.1 for SQK-LSK110 libraries, R10.4.1 for SQK-LSK114 libraries) and sequenced on an ONT PromethION P48 device. Samples were run for a maximum duration of 72 hours, with 1-3 nuclease flushes and reloads performed during the run, where necessary to maximise sequencing yield.

### ONT data processing

Raw ONT sequencing data was converted from FAST5 to the more compact BLOW5 format^36^ in real-time on the PromethION during each sequencing run using *slow5tools*^*37*^ (v0.3.0). BLOW5 data was transferred to the Australian National Computational Infrastructure (NCI) high-performance computing environment before further processing. Data was base-called using Guppy (v6.0.1) with the high-accuracy model and reads with mean quality < 7 were excluded from further analysis.

### Alignment to reference genome

To evaluate the use of *hg38* and *T2T-chm13* reference genomes, ONT libraries generated in our study for the HG001 and HG002 reference samples and matched Illumina libraries from the GIAB consortium were mapped against each reference genome. The short read data was mapped using *bwa-mem2* (v2.2.1), with *-Y* optional parameter, and the long read data was mapped using *minimap2*^38^ (v2.22) with the following optional parameters: *-x map-ont -a --secondary=no --MD*. The alignment of each individual library to either *hg38* or *T2T-chm13* was made in a sex-specific manner with an XY reference for genotypically male individuals and an XO reference for genotypically female individuals. After selecting *T2T-chm13* as our central reference genome, all other ONT libraries in the cohort were also mapped to this reference using *minimap2*, as just described.

### Detection of structural variation

Detection of large indels (20-49bp) and SVs (≥ 50bp) on HG001 & HG002 Illumina mapped libraries was performed using *smoove* (v0.2.6) with default parameters. Variant detection with ONT mapped libraries was performed on each individual sample using *CuteSV*^15^ (v1.0.13) with the following optional parameters: *--max_cluster_bias_INS 100 --diff_ratio_merging_INS 0*.*3 --max_cluster_bias_DEL 100 --diff_ratio_merging_DEL 0*.*3 --report_readid -- min_support 5 --min_size 20 --max_size 1000000 --genotype*. Deletions with < 20% of supporting reads for the variant sequence were excluded and insertions with < 5% of supporting reads were also excluded. Individual callsets were then merged into a unified joint-call catalogue using *Jasmine*^18^ with the following optional parameters: *min_support=1 --mark_specific spec_reads=7 spec_len=20 --pre_normalize --output_genotypes -- allow_intrasample --clique_merging --dup_to_ins --normalize_type --run_iris iris_args=min_ins_length=20,-- rerunracon,--keep_long_variants*. Variants in the joint non-redundant callset were filtered to exclude events with weak evidence (QUAL ≤ 5).

### Structural variation repeat classification

Indels and SVs were classified according to repeat type using custom analysis methods. We first created an extended local allele sequence for each variant, which was 5x the size of the variant itself. For each insertion, we created an extended ALT allele by extracting reference sequence from the *T2T-chm13* genome immediately upstream (2x variant size) and downstream (2x variant size) of the variant site, then concatenating these in appropriate order with the consensus insertion sequence that was retrieved from the Jasmine VCF. For each deletion, we created an extended REF allele by extending the variant position in either direction (by 2x variant size) and extracting reference sequence from the *T2T-chm13* genome.

Each extended allele, which captures the variant in its local sequence context, was then scanned for tandem repeats using *Tandem Repeat Finder*^39^ (trf409.linux64) with input parameters recommended by the developers (*2 7 7 80 10 50 500*). Annotated tandem repeats were parsed by their period: 1bp=homopolymer (HOMO); 2-12bp=short-tandem-repeat (STR); >12bp=tandem-repeat (TR). Any overlapping annotations of the same type were merged. We then calculated the extent to which the variant site (i.e. central 20% of the local sequence allele) was covered by repeats of each type; if ≥75% of the variant was covered by repeats of a single type, the variant was classified accordingly as either ‘HOMO’, ‘STR’ or ‘TR’.

Each extended allele was then scanned for interspersed mobile elements using *RepeatMasker* (4.1.2-p1) with the following input parameters: *-species human -gff -s -norna -nolow*. Annotated interspersed repeats were parsed into different types (SINE, LINE, DNA Transposon, LTR, Retroposon, Other), based on RepeatMasker classifications, and labelled as ‘Complete’ (≥75%) or ‘Fragment’ (<75%) based on the fraction of the canonical sequence element that was present. We then calculated the extent to which the variant site within its local allele sequence was covered by interspersed repeats; if ≥75% of the variant was covered by an element or elements of a single type, the variant was classified accordingly as either: ‘SINE’, ‘LINE’, ‘DNA Transposon’, ‘LTR’, ‘Retroposon’ or ‘Other’. If the variant itself covered at least ≥75% of one or more complete annotated elements, the variant was labelled as a ‘Complete’ transposition event. If not, it was labelled as a mobile element ‘Fragment’, which are mostly small SVs contained within larger interspersed elements. Variants that were not classified with either a tandem or interspersed repeat label, were considered ‘Non-repetitive’.

### Comparison to annotations

To assess the novelty of our SV catalogue, we compared SVs to: (*i*) the *gnomAD* (v2.1) SV database (ncbi.nlm.nih.gov/sites/dbvarapp/studies/nstd166/) and; (*ii*) an SV callset from population-scale ONT sequencing of Icelanders published recently by deCODE genetics (github.com/DecodeGenetics/LRS_SV_sets). For this analysis, we first converted SV coordinates from *T2T-chm13* to the *hg38* reference genome using the *LiftOver* utility from UCSC. SVs that were successfully lifted over were then intersected with the *gnomAD* and *deCODE* annotations, separately. To account for variability in methods of SV detection between the different studies, as well as potential confounding effects during *LiftOver*, we allowed for some discrepancy in the placement of breakpoints; to be classified as ‘Found’, an SV had to share ≥80% reciprocal similarity with an intersecting SV in one or both of the *gnomAD* and *deCODE* annotations.

### Telomere distance

The distance of each variant to the nearest telomere was calculated and based on that distance variants were binned into 500 kb fixed windows. The number of variants in each bin was counted and averaged across all chromosomes to assess the density of variant distribution across a generic chromosome. There was a higher density of variants within the 5 Mb of the telomere region, including acrocentric and metacentric chromosomes. To investigate the types of variants driving this effect, we parsed the variant counts in the 500 kb bins based on variant type. We then fitted a LOESS curve onto the log-transformed counts of the different bins for each variant type, displaying the results in an exponential scale, to ensure that y-values in the regression were all greater than 0. The curves for different variant types showed that tandem repeats were the variants mostly driving the higher density near telomeres.

### Principal Coordinate Analysis

The non-redundant variant callset was converted into a matrix, where each row represented a variant and each column represented an individual. The presence of a variant in an individual was represented with a 1 and the absence with a 0. We then used the *vegdist* function from the *vegan* R package to calculate the dissimilarity between individuals based on their variant composition using the Bray-Curtis method. From the dissimilarity index matrix, we performed a Principal Coordinate Analysis (PCOA) using the *pcoa* function from the *ape* R package, plotting the Principal Coordinate Axis 1 & 2 (PCoA1 & PCoA2) for each individual according to their community/group.

### Discovery curves

To measure SV diversity, we generated discovery curves, wherein we calculate the number of new non-redundant SVs gained as additional individuals are considered. Starting with a single NCIG individual, the number of non-redundant variants was calculated each time a new individual was added to the analysis, until all 141 NCIG individuals were included. The growth rate of the nonredundant set declines as the number of cumulative individuals increases. We then used the values obtained in the discovery curve to generate a log regression model of the number of non-redundant variants as a function of the number of individuals sampled with the *lm* function from the *stats* R package. The curves model the level of heterogeneity in a given group, and enable estimation of the number of individuals required to saturate variant discovery. We then generated discovery curves and log regression models by parsing the variants for each community/group (NCIGP1, P2, P3, P4, non-NCIG and NCIGP1/P2/P3 combined), according to geographical distribution (NCIG-only, NCIG-absent, Global) and variant type (Non-repetitive, Tandem repeats & Mobile elements).

### LOEUF constraint analysis

We binned variants in the non-redundant callset intersecting protein coding genes with the LOEUF decile of that corresponding gene previously assigned by *gnomAD*, and which measures intolerance to loss-of-function variation. Genes in the 1^st^ decile have the highest constraint, while genes in the 10^th^ decile are the least constrained. Therefore, if variants (large indels and/or SVs) regularly have deleterious effects on gene function, the expectation would be that genes in the 1^st^ decile would harbour relatively fewer variants than genes in higher deciles, after accounting for gene size. To test this, we calculated the variant density within CDS and non-CDS regions (introns, UTRs & +/-2 kb flanking regulatory regions) of all the genes in each LOEUF decile. Density was calculated by counting the number of non-redundant variants intersecting CDS regions of all genes in a given decile, divided by the total size of CDS regions of all genes of that decile. Similarly, we counted the number of variants intersecting non-CDS regions (but not intersecting CDS regions) and divided that by the total size of non-CDS regions of all the genes in a given decile. Variant density was plotted per LOEUF decile for CDS & non-CDS regions, showing clear differences between high and low deciles for both CDS and non-CDS regions.

### Analysis of short-tandem repeats

To explore the landscape of STR variation, we retrieved all joint-called indels and SVs classified above as ‘STR’ variants, which represent expansions (i.e. insertions) and contractions (i.e. deletions) of local STR elements. For each variant, we recorded the total expansion/contraction size, the period size (2-12bp) and the STR motif identified by *Tandem Repeat Finder* (see above), and investigated global frequencies for each of these dimensions. For STR motif frequency analysis, we considered all possible motif representations in both orientations as a single redundant motif (e.g. CAG, AGC, GCA, TGC, GCT, CTG are a single redundant triplet). To identify significantly expanded STR sites, we applied the following criteria: (i) STR period size ≥3bp; (ii) local STR element expanded by at least ≥10 repeats; (iii) local STR element is expanded by ≥50% of its reference size; (iv) local STR element reference size is <1kb. This identified 651 STR sites within protein-coding genes that were significantly expanded in at least one individual.

Individual-level, diploid genotyping of STR alleles was used to elucidate full allelic diversity at the 651 STR sites within protein-coding genes that were significantly expanded in at least one individual (see above), as well as 50 known disease-associated STR loci. This was performed using a custom analysis method. Briefly, we identified variation within the local region around a given STR site using *clair3*^40^ (v0.1-r12; for SNVs and 2-20bp indels) and *sniffles2*^41^ (v2.0.2; for indels/SVs > 20bp). *Sniffles2* was used instead of *CuteSV* (as above) because it shows better performance at STR sites when guided by the *--tandem-repeats* input parameter. Variants from *clair3* and *sniffles2* were incorporated in a haplotype-specific fashion into the local genome sequence using *bcftools consensus* (v1.12)^42^, and the modified hap1/hap2 sequences were extracted in a +/-50bp window centred on the STR site; these constitute the consensus STR allele sequences for a given individual at a given STR site, with the larger being designated ‘allele_A’ and the shorter ‘allele_B’. *Tandem Repeat Finder* was used to determine the STR period size, length, motif and other summary statistics for each STR allele. Allele sequences were visualised in sequence bar charts, in which each tile represented a nucleotide (A, C, G & T), using *R* package *ggplot2*. To investigate the variability of STRs within the different communities, we calculated the mean and standard deviation of STR lengths for the alleles within each community. We then performed an ANOVA test (p-value < 0.05) to identify STR sites that were significantly variable between communities. We plotted the standard deviation for each significantly variable site, normalised to range between 0 and 1, as a heatmap and also performed hierarchical clustering using the *heatmap*.*2* function of the *gplots* R package.

### Data analysis

All data manipulation and visualisation, as well as plotting was performed in R (v4.0.0).

## DATA AVAILABILITY

Requests for materials and data should be addressed to the NCIG Data Access Committee, John Curtin School of Medical Research, Australian National University

## CODE AVAILABILITY

All code used in the analysis and figure generation will be shared publicly at the time of publication.

## ACKNOWLEDGEMENTS

We are indebted to the individuals and their communities who participated in this research and to the NCIG Governance Board who helped guide this work in a culturally appropriate manner. We thank our colleague Daniel MacArthur for helpful feedback during manuscript preparation. This project was undertaken with the assistance of resources and services from the National Computational Infrastructure (NCI), which is supported by the Australian Government and the Australian National University (ANU). We acknowledge the following facilities that were used during this study: the Garvan Institute Sequencing Platform (Nanopore sequencing service) and the Australian Phenomics Facility (APF). We acknowledge the following funding sources: Medical Research Futures Fund grants MRF2016008, MRF1173594 & MRF2016124, National Health and Medical Research Council grants 2011277 & 2021172, and philanthropic support from The Kinghorn Foundation.

## DECLARATIONS

This project receives partial in-kind support from Oxford Nanopore Technologies (ONT) under an ongoing collaboration agreement. A.R., J.M.H., H.G. & H.P. have previously received travel and accommodation expenses from ONT to speak at conferences. H.G. and I.W.D. have paid consultant roles with Sequin PTY LTD. The authors declare no other competing financial or non-financial interests.

## CONTRIBUTIONS

I.W.D. and H.R.P. conceived the study, with the support of G.J.M. and A.H. The NCIG provided access to DNA samples from Indigenous communities. O.M.S. and A.W.H. provided non-Indigenous comparison samples. M.R., J.M.H., I.S., M.A.K. and D.S.B.D. processed samples and performed sequencing experiments. A.L.M.R., H.G., S.R.C., H.R.P. and I.W.D. performed bioinformatics analysis. A.L.M.R. and I.W.D. generated the Figures. A.L.M.R. and I.W.D. wrote the manuscript, with input from B.L., A.B., G.B., G.J.M., A.H. and H.R.P.

