## Extended Data Figures for "The landscape of genomic structural variation in Indigenous Australians"

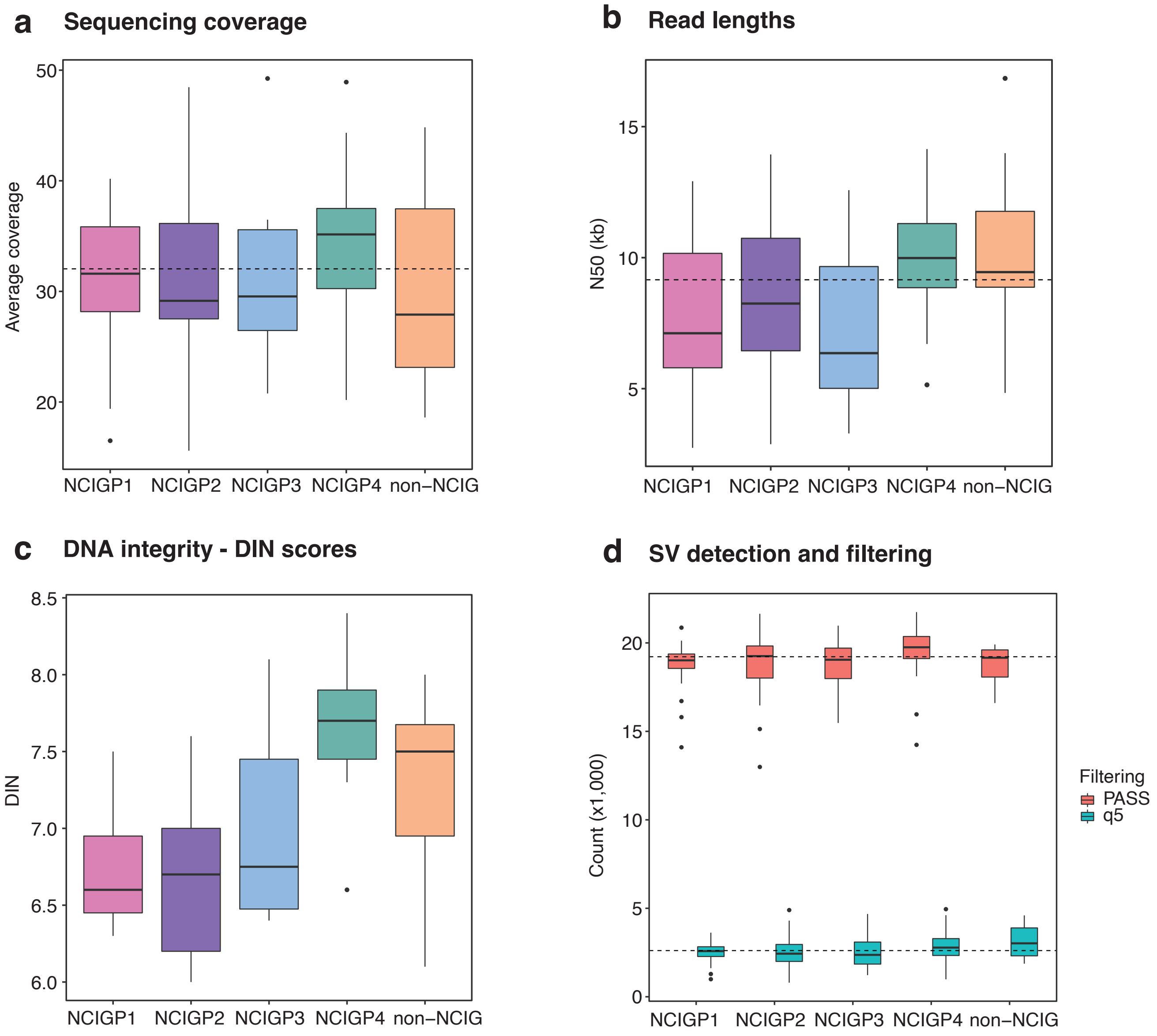


**Extended Data Fig1. (a)** Boxplot shows the average depth of coverage per individual grouped by their communities (NCIGP1 = pink, P2 = purple, P3 = blue, P4 = green & non-NCIG = orange). The horizontal dashed line indicates the average coverage across all libraries in the cohort. (**b**) Boxplot shows the N50 distribution of individual libraries in the different communities. The horizontal dashed line indicates the average N50 across all libraries in the cohort. **(c)** Boxplot shows the distribution of DNA Integrity Number (DIN), which indicates the level of fragmentation of a genomic DNA sample, for individual libraries across the different communities. **(d)** Boxplot shows the distribution of the number of high-quality (PASS=orange) and low-quality (q5=green) structural variants per individual grouped by community after quality filtering (Quality ≥ 5). The horizontal dashed lines indicate the average number of high-quality (top) and low-quality (bottom) structural variants across all libraries in the cohort.

**Repetitive medically-relevant genes, such as *MUC1*, are best resolved by long-read sequencing aligned to the *chm13-T2T* reference.**

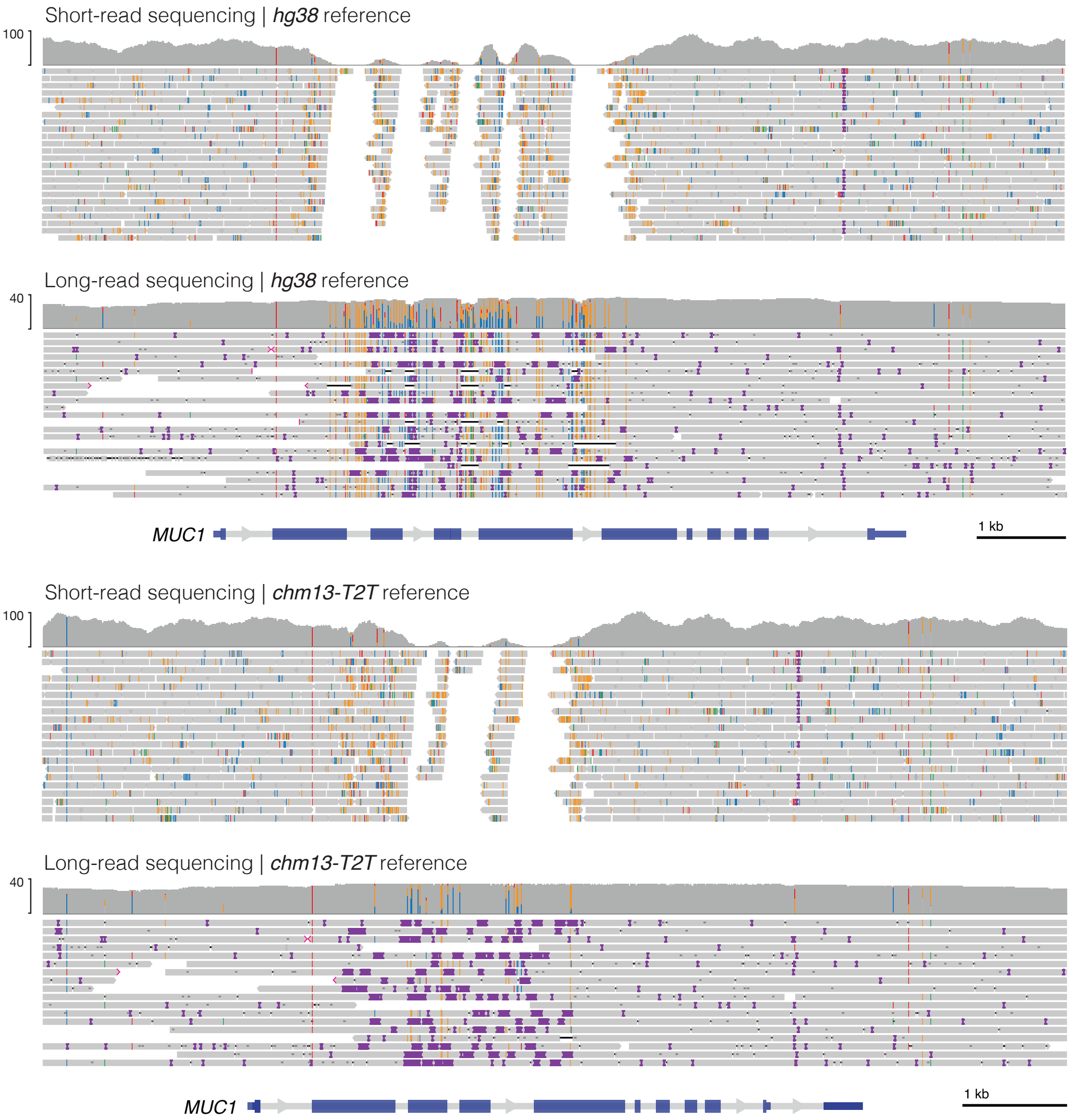


**Extended Data Fig2.** Genome browser views show comparison of short-read and long-read alignments to either the hg38 or chm13-T2T reference genomes at MUC1, an example of a repetitive medically relevant gene. Both datasets are from the HG002 reference sample. The gene contains a large tandem repeat region that is best resolved by alignment of long-reads to chm13-T2T.

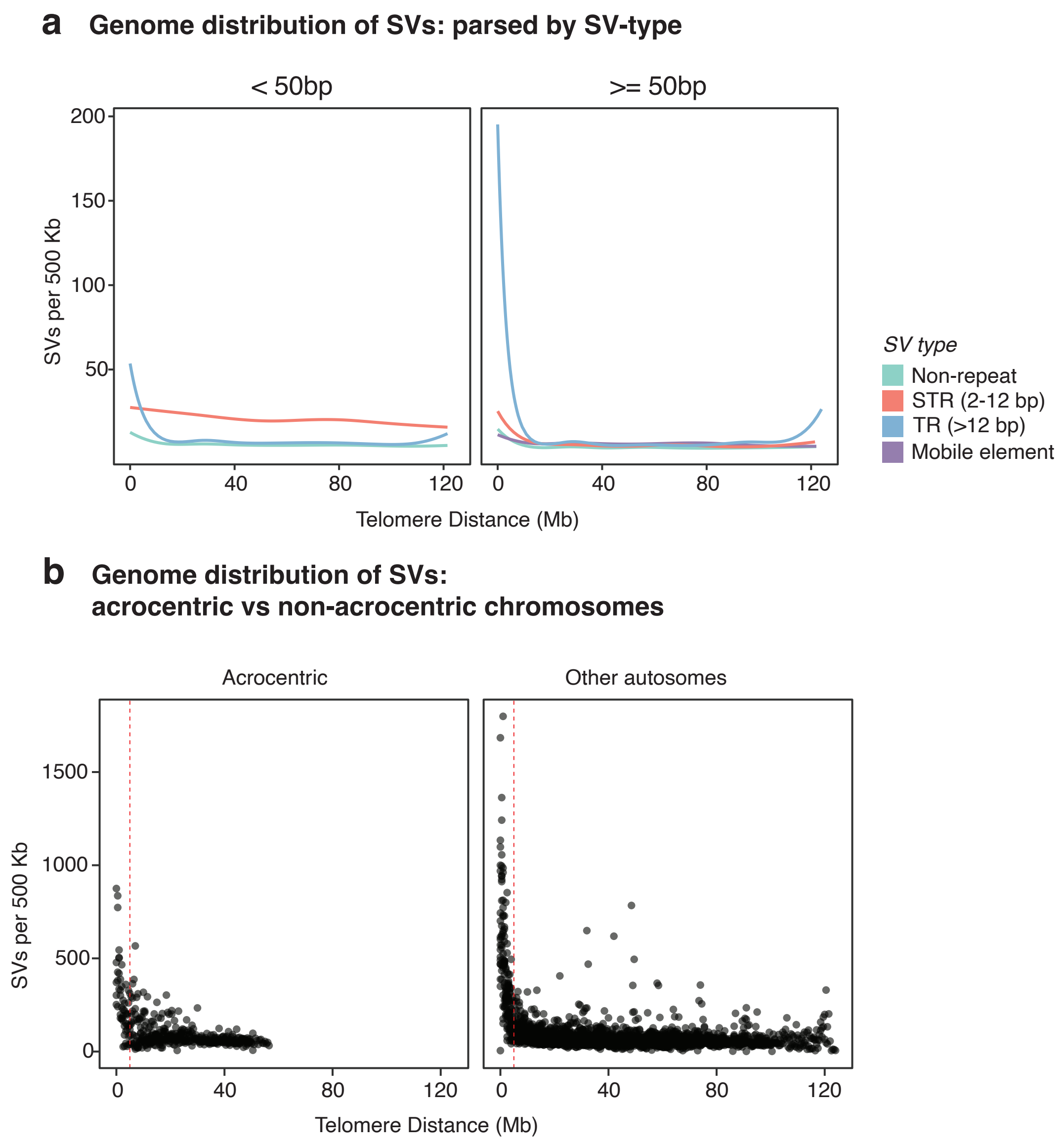


**Extended Data Fig3. (a)** Line plots show LOESS curves of the number of large indels (> 20 & < 50 bp) and structural variants (≥ 50 bp) per 500 Kb fixed window relative to the distance to the nearest telomere, parsed by variant type (non-repetitive = 0, short tandem repeat = red, tandem repeat = blue & mobile element = purple). **(b)** Dot plots show the number of structural variants per 500 Kb fixed window, relative to the distance of the window to the nearest telomere, for acrocentric and metacentric autosomes. The vertical dashed lines indicate a distance of 5 MB from the telomere.

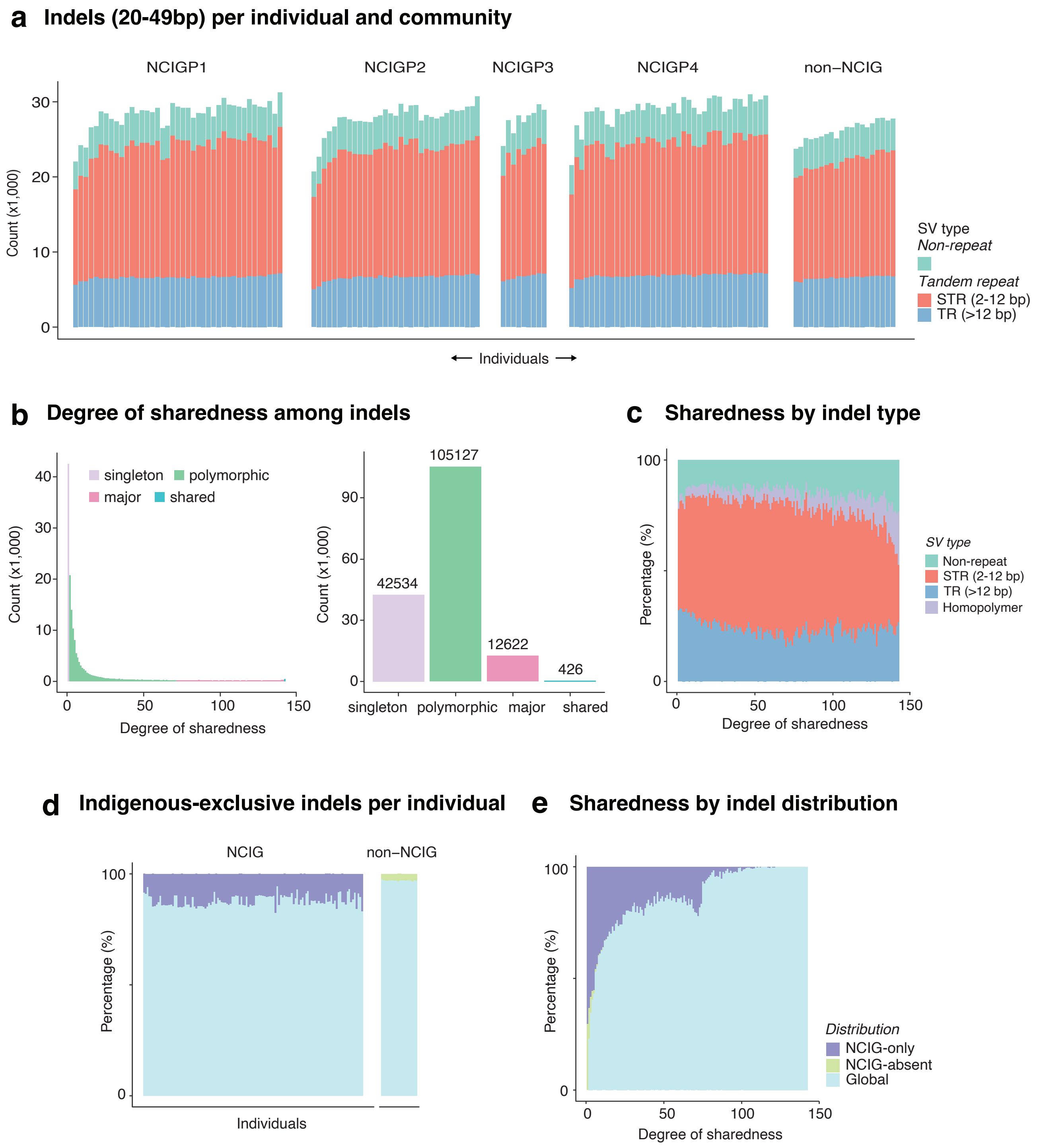


**Extended Data Fig4. (a)** Bar chart shows the number of large indels (20-49 bp) identified in individuals from each group (NCIGP1, P2, P3, P4 & non-NCIG), broken down by type: non-repeat (green) and tandem repeat (STR = red & TR = blue). **(b)** Bar plot shows the number of non-redundant structural variants identified within a given number of individuals in the cohort (degree of sharedness). Variants were classified as private (1 individual), polymorphic (> 2 & < 50% of individuals), major (≥ 50% of individuals & < all individuals) and shared (all individuals). **(c)** Bar chart shows the proportion of different variant types (same colour scheme as a, in addition to homopolymers = light purple) for large indels identified within a given number of individuals in the cohort (degree of sharedness). **(d)** Bar chart shows the proportion of large indels in each individual that were only found in NCIG individuals (NCIG-only = purple), only found in non-NCIG individuals (NCIG-absent = green) and found in both NCIG & non-NCIG individuals (Global = light blue). **(e)** Bar chart shows the proportion of NCIG-only, NCIG-absent and Global large indels (same colour scheme as d) for all the variants identified within a given number of individuals in the cohort (degree of sharedness).

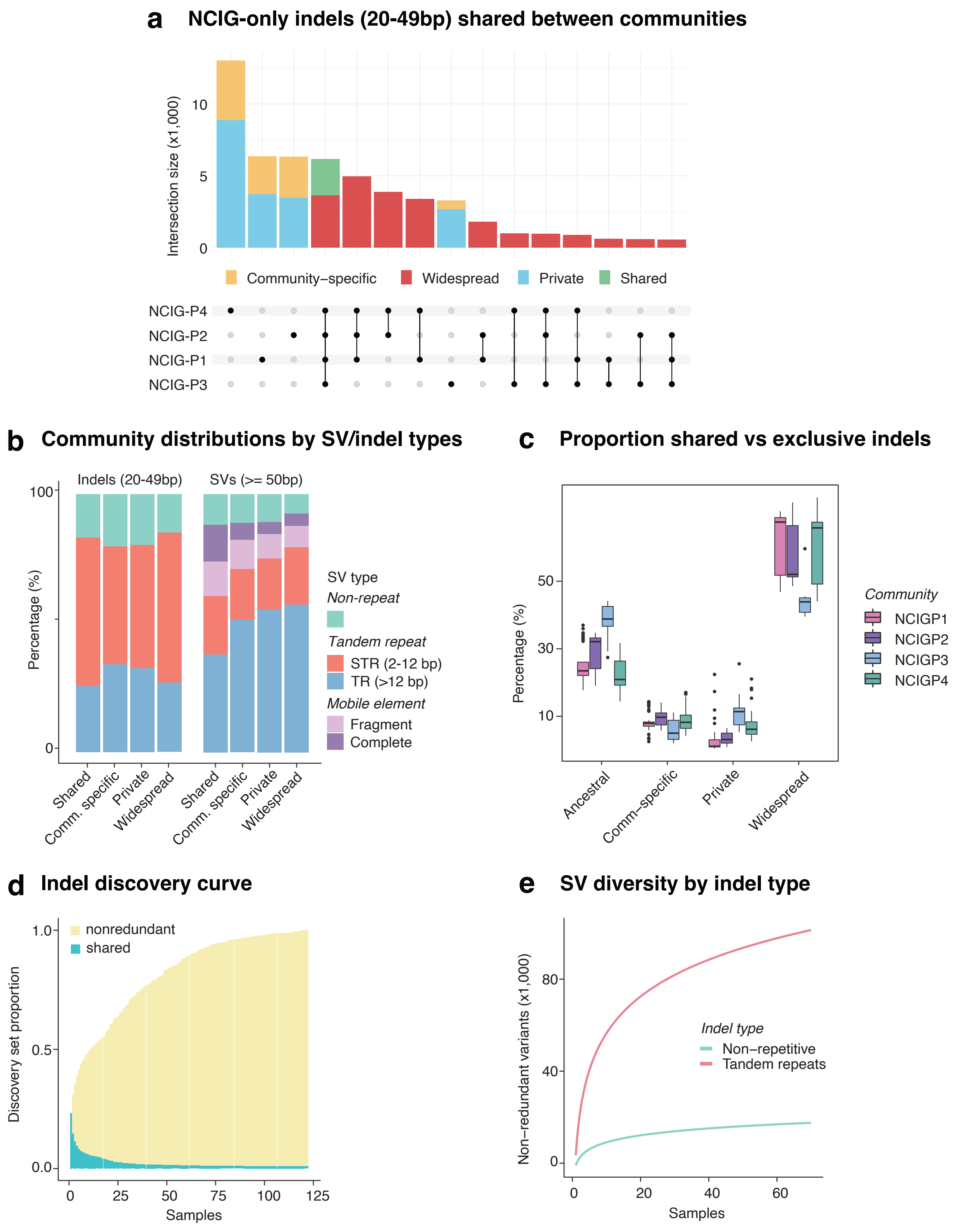


**Extended Data Fig5. (a)** Upset plot shows the distribution of NCIG-only large indels (20-49 bp) shared among the four indigenous communities (NCIGP1, P2, P3 & P4). Variants were classified as private (n = 1 individual; blue), community-specific (n > 1 individual in 1 community; yellow), widespread (n > 1 individual in more than 1 community; red) or shared (n > 1 individual in all 4 communities; green) according to the number of communities in which they were identified. **(b)** Proportion of different SV types for NCIG-only variants classified as private, community-specific, widespread or shared. Types are non-repetitive (teal), tandem repeat (STR = red & TR = blue) and mobile element (fragment = light purple & complete = dark purple). **(c)** Proportion of private, community-specific, widespread & shared NCIG-only variants among individuals, grouped by community. **(d)** Bar chart shows a discovery curve, in which starting with a single NCIG individual, the number of new non-redundant large indels is counted by iteratively adding the unique calls from additional NCIG individuals. Indels shared among all previously added samples are shown as green portions of each bar. The growth rate of the nonredundant set declines as the number of samples increases. **(e)** Log regression model showing the predicted number of non-redundant large indels identified given the number of individuals sampled. The model was broken down by variant type (Non-repetitive = green, Tandem repeats = red).

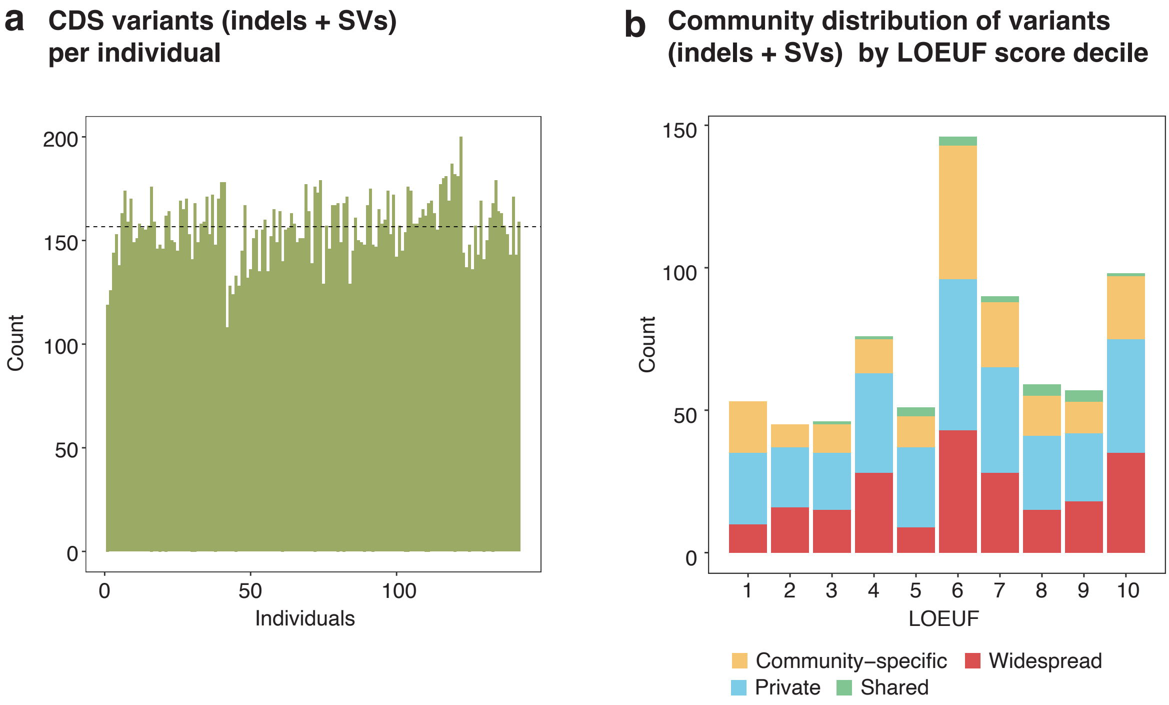


**Extended Data Fig6. (a)** Bar plots show the number of variants per individual impacting CDS exons of protein-coding genes. The horizontal dashed lines indicate the average number of variants in CDS regions across the entire cohort. **(b)** The bar plot shows the number of NCIG-only variants per LOEUF decile parsed by their level of distribution within NCIG communities. Variants were classified as private (n = 1 individual; blue), community-specific (n > 1 individual in 1 community; yellow), widespread (n > 1 individual in more than 1 community; red) or shared (n > 1 individual in all 4 communities; green) according to the number of communities in which they were identified.

**a** **Identifying STR sites in protein coding genes with significant expansions**

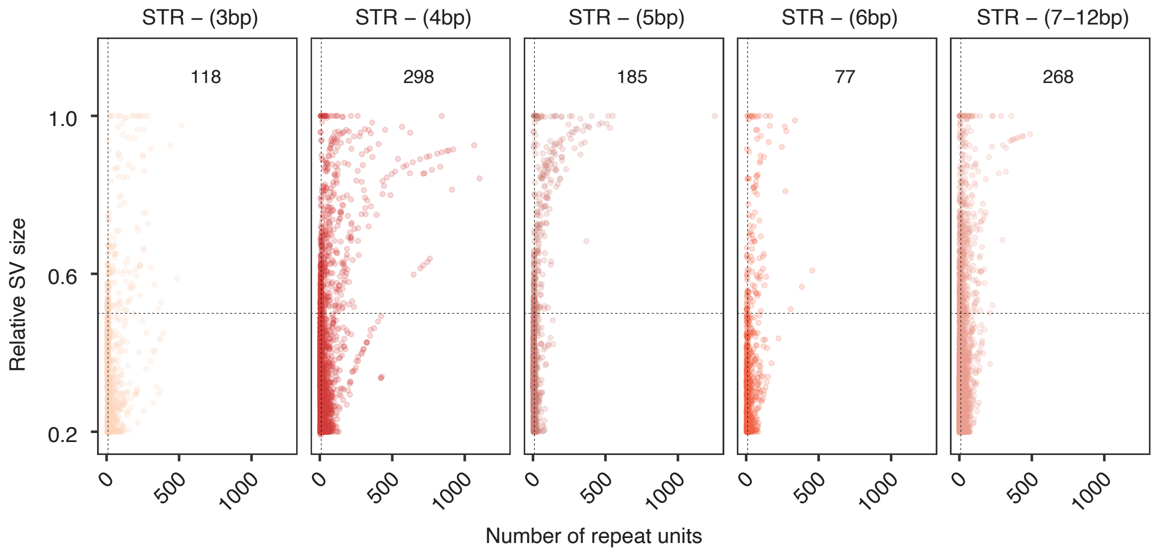


**b** **Diversity of STR expansions within protein-coding genes**

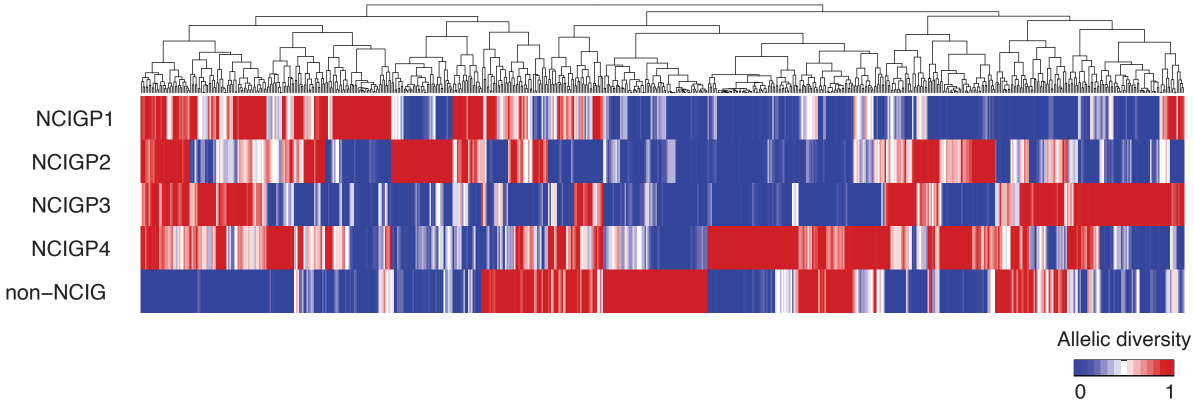


**Extended Data Fig7. (a)** Dot plots show the number of repeat units versus the relative size increase of STR expansions of different period sizes. The horizontal dashed line indicates the minimum relative size increase (0.5) and the vertical dashed line indicates the minimum number of repeat units (10) required for an STR expansion to be further genotyped across all individuals in the cohort. **(b)** Matrix shows the normalised standard deviation (range 0:1) of allele sizes within each community for all STR sites with expansions in one or more individuals, in which allelic composition between the groups was significantly different. Hierarchical clustering was performed to group the STR sites based on the different patterns of variability between communities.
